# ActSeek: Fast and accurate search algorithm of active sites in Alphafold database

**DOI:** 10.1101/2025.02.11.637678

**Authors:** Sandra Castillo, O. H. Samuli Ollila

## Abstract

Finding proteins with specific functions by mining modern databases can potentially lead to substantial advancements in wide range of fields, from medicine and biotechnology to material science. Currently available algorithms enable mining of proteins based on their sequence or structure. However, activities of many proteins, such as enzymes and drug targets, are dictated by active site residues and their surroundings rather than the overall structure or sequence of a protein. Here we present ActSeek – a computer vision-inspired fast program – that searches structural databases for proteins with active sites similar to the seed protein. ActSeek is implemented to mine proteins with desired active site environments from the Alphafold database. The potential of ActSeek to find innovative solutions to the world’s most pressing challenges is demonstrated by finding enzymes that may be used to produce biodegradable plastics or degrade plastics, as well as potential off-targets for common drug molecules.

## Introduction

Finding proteins with desired functions by mining databases is becoming increasingly attractive due to the growing amount data and emerging computational tools. In addition to Uniprot database (1) containing protein sequences, also *in silico* predicted protein structures are available for mining (2, 3). These databases open new avenues for searching proteins based also on structural similarity, in addition to traditional sequence based approaches using BLAST or similar algorithms (4). Such algorithms are available, but their computational efficiency is often a limiting factor for large scale structure based searches (5–8).

A significant improvement in performance was presented in the recent FoldSeek algorithm (7), which enabled structure-based searches over a large number of structures available in databases. However, the activities of some relevant proteins, such as enzymes and drug targets, is dictated by several residues in the proximity of active sites rather than the overall structure of a protein. Therefore, searches based on full structure may result in proteins whose overall structure resembles the seed protein, but have critical differences at the active site region. On the other hand, proteins with similar active site to the seed protein but some irrelevant structural differences may not be found.

Here we present ActSeek, a program that searches structural databases for proteins with similar active site to the seed protein. Our fast algorithm, inspired by computer vision (9, 10), enables searches over large databases based on the coordinates of specific amino acids. Here we have implemented ActSeek to search over the Alphafold database (3).

Finding proteins with desired active sites and functions can benefit wide range of fields, from medicine and biotechnology to material science. Here we demonstrate the potential of ActSeek to find (i) proteins that are able to produce biodegradable plastics with enhanced properties (polyhydroxyalkanoate (PHA) synthases) (11, 12), (ii) new proteins to degrade poly(ethylene terephthalate) (PET) based plastics (depolymerases PETase and MHETase) bearing potential in recycling applications (13), and (iii) potential off-targets for common drugs. These examples demonstrate the potential of protein mining based on active site structures in providing innovative solutions to the world’s most pressing challenges.

While other methods can find structural motifs with high speed as ActSeek does (14–16), they are limited to smaller databases such as the RCSB PDB (17) or BRENDA (18). This limitation makes it difficult to search larger or customized protein databases. ActSeek, on the other hand, searches in the AlphaFold database (3), which contains over 200 million proteins, or any protein set the user provides.

## Results and Discussion

### A. ActSeek algorithm

In contrast to BLAST (4) and Fold-Seek (7) that are based on overall sequence or structure of proteins, ActSeek finds proteins with similar local structure to the active site of a seed protein (Fig. 1A). Here we implement the search from the Alphafold database (3), yet the algorithm can be applied to other databases with 3D structures of proteins under interest.

**Fig. 1.**
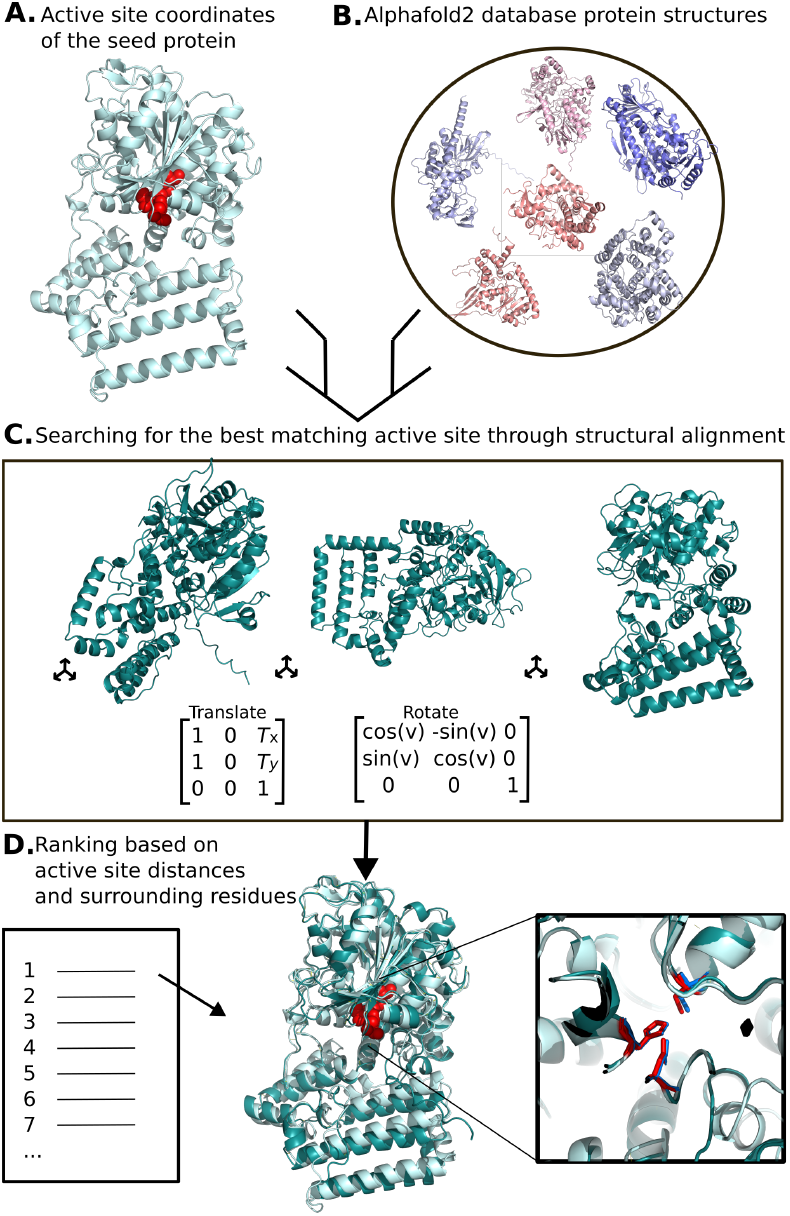
**A** Structure of an example seed protein with active site amino acids marked with red. **B** Query proteins are taken from a data (alphafold database (3) in this work). **C** Optimal rotational and translational matrices found with SVD and least square fitting (10). **D** Best targets selected based on local structure.

To initiate the ActSeek search, user needs to first provide the structure of the seed protein and define the relevant amino acids for the active site and its surroundings (Fig. 1A). Alternatively, the user can provide the coordinates of the alpha and beta carbons of the relevant amino acids. Three most relevant amino acids defined by the user, corresponding for example catalytic or binding site, will be used to compute the optimal rotational and translational matrices for structural alignment of the seed and query proteins using singular value decomposition (SVD) and least square fitting (10) (Fig. 1C). After euclidean transformation using these matrices, query proteins found from a given database are ranked based on distances between amino acids in the seed and the query structures. These distances are calculated using binding site amino acids together with additional ones defined by the user, and their neighboring residues. As a results, ActSeek provides a ranked list of natural enzymes with similar 3D arrangement of the active site residues to the seed (Fig. 1C).

To maximize the speed of the algorithm and enable searches over large datasets, query sequences are pre-screened before proceeding to computationally expensive comparison of 3D arrangements between seed and query proteins. This is done by checking that a query structure contains the three active site amino acids with mutual distances similar to the seed.

### B. Searching new enzymes

To demonstrate the utility of ActSeek for finding new enzymes, we searched proteins that contain active sites characteristic for PHA synthases (19), and two different PET depolymerases PETase and MHETase (20). PHA synthases are used to produce biodegradable plastics (11, 12), while PETase and MHETase degrade widely used plastics bearing potential in recycling applications (13, 21). ActSeek results for these enzymes are compared with the sequence based BLAST (4) and structure based FoldSeek (7) in Fig. 2 a.

**Fig. 2.**
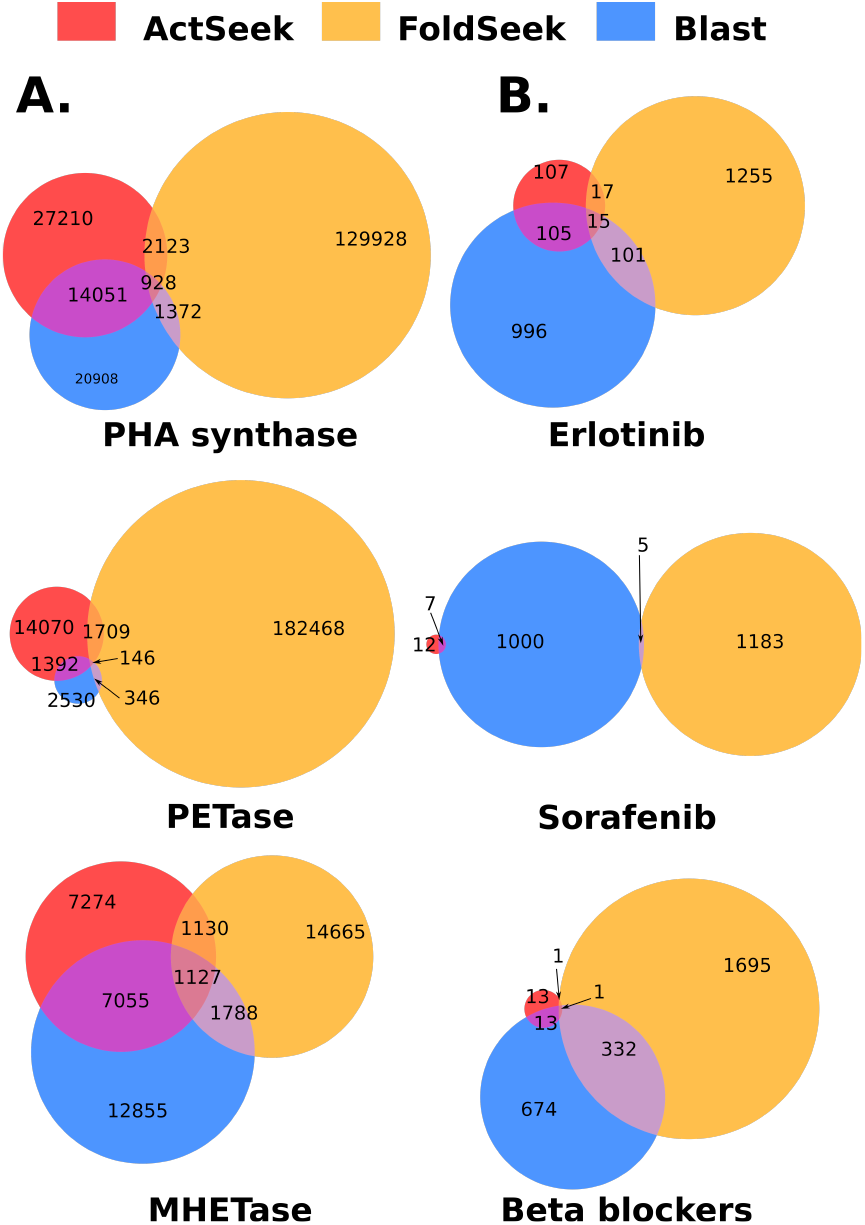
**A** Results for ActSeek searches over proteins in all organisms using active sites of a PHA synthase, PETase and MHETase as seeds. **B** Results for ActSeek searches over human proteins with relevant binding pocket amino acids for eriotinib, sorafenib and beta blockers as seeds.

For all these enzymes, ActSeek finds significant amounts of proteins that are not recognized by BLAST or FoldSeek searches (11964 for PHA synthase, 11115 for PETase and 216 for MHETase). This means that large number of enzymes containing the relevant active site, thereby potentially having the desired functions, are not found by BLAST or FoldSeek searches. On the other hand, these methods find many proteins with similar sequences or overall structures to the seed, but that do not contain the active site required for the desired reaction. These results demonstrate that ActSeek can 1) find new proteins with the desired active site that are not found by searches based on sequences or overall structures, and 2) filter out proteins that do not contain the relevant active sites from BLAST or FoldSeek results. The new natural enzymes found by ActSeek can bear novel properties that are potentially useful in wide range of applications from specificity for relevant substrates to functionality in harsh conditions.

Furthermore, our results demonstrate the potential of ActSeek to improve functional annotation of proteins. For example, 31585 proteins are currently annotated as PHA synthases in the Uniprot database (1). However, 9080 of these do not contain the relevant active site according to the ActSeek. On the other hand, ActSeek finds 4875 proteins with the relevant active site for PHA synthesis, but that are not annotated for this functions in Uniprot database. Based on these results, ActSeek can provide new annotations for protein functions based on active sites, as well as filter incorrect annotations from databases.

### C. Searching off-target proteins for drugs

Beside enzymes, drug binding to query proteins is another example where local binding region is more relevant than the overall structure or sequence. Here we demonstrate that ActSeek can find potential off-target proteins for drug molecules. To this end, we searched all human proteins from Alphafold database with binding pockets for two anti-cancer drugs, Erlotinib (22) and sorafenib (23), and for beta-blockers used to treat various cardiovascular diseases such as hypertension, cardiac arrhythmia, or myocardial infarction and also alleviate severe anxiety (24). Erlotinib and sorafenib inhibit epidermal growth factors and protein kinases, respectively, and beta-blockers block receptor sites of adrenergic receptors. ActSeek results using the main targets of these drugs as seeds (Uniprot P00533, P15056, P07550, respectively) are compared with the sequence based BLAST (4) and structure based FoldSeek (7) in Fig. 2 b.

ActSeek finds relatively few proteins with suitable active sites for binding of drugs studied here (107 for Erlotinib, 12 for Sorafenib and 13 for beta blocker). Majority of ActSeek results are found also by Blast search based on sequences, but Blast finds also hundreds of proteins that do not contain the relevant binding site according to ActSeek, therefore being unlikely targets for the given drug (Fig. 2b). Structure based FoldSeek search finds hundreds of proteins with similar structure to the seed, but only few of them (17 for Erlotinib and one for beta-blocker off-targets) contain the relevant binding site, compromising its suitability for finding drug off-targets.

For Erlotinib, ActSeek results contain two proteins that were not identified with sequence based Blast search (Fig. 2). These two proteins have less than 30% sequence similarity with the seed protein and only partial structural similarity, but they have relevant amino acids in the exact locations as in the seed protein (Fig. 3), which explains why they were found by ActSeek but not by Blast or FoldSeek. Previously reported off-targets for Erlotinib that may contribute to the common side-effects (25, 26) were identified by both ActSeek and Blast, but not by FoldSeek.

**Fig. 3.**
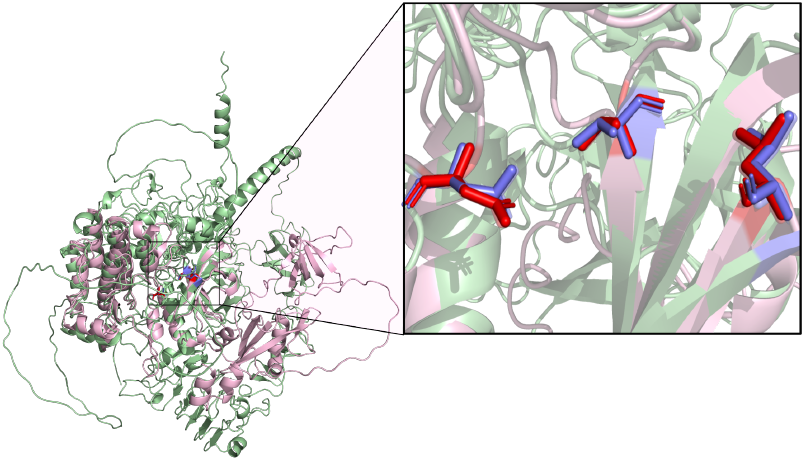
Potential off-target for Erlotinib, tyrosine-protein kinase Srms (Uniprot: Q9H3Y6) (pink) superposed to the seed protein epidermal growth factor receptor (Uniprot: P00533) (light green). This protein was found by ActSeek but not by Blast or FoldSeek. The amino acids used for the search of Erlotinib off-targets are marked as blue (718Leu, 790Thr and 855Asp). The corresponding amino acids in the tyrosine-protein kinase Srms are marked as red (236Leu, 302Thr and 368Asp).

For Sorafenib, we identified 11 potential off-targets. These include the retinal guanylyl cyclase2 (Uniprot P51841) found also by Blast, which may explain one of the reported drug’s side effects related to the retinal tear (27). Additionally, two of the identified drug targets are annotated as atrial natriuretic peptide receptor 1 and 2 (Uniprot P16066 and P20594) and were found only by ActSeek. Alignment of atrial natriuretic peptide receptor 1 with the main Sorafenib target, B-Raf kinase (Uniprot P15056), with Sorafenib docked is shown in Fig. 4. Inhibiting these receptors may cause high blood pressure and heart-related problems, which are also reported side effects of the drug (28). However, none of these receptors have been previously reported as off-targets of the drug. Also, previously reported off-targets for Sorafenib, VEGFR 1, 2 and 3 (29), are among the ActSeek results.

**Fig. 4.**
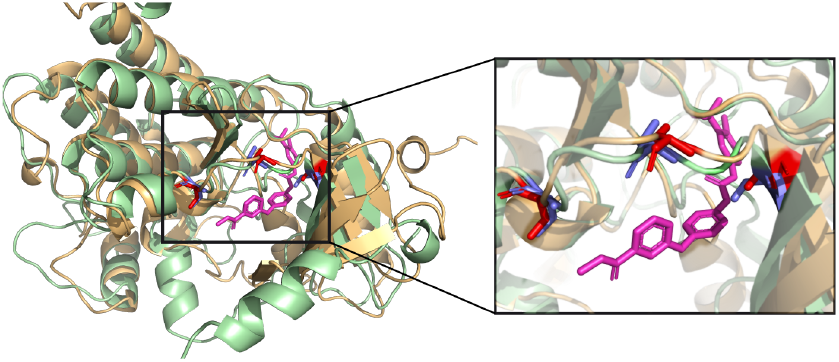
Potential off-target for Sorafenib, atrial natriuretic peptide receptor 1 (Uniprot P16066, beige), aligned with the main target, B-Raf kinase (Uniprot P15056, light green). Amino acids involved in the binding are marked as red in the B-Raf kinase and blue in the atrial natriuretic peptide receptor 1. Sorafenib (magenta) has been docked to the B-Raf kinase structure.

All 12 potential off-targets for beta-blockers found by Act-Seek are identified also by Blast. Successfully recognized case by Blast with relevant active site, 5-hydroxytryptamine receptor, is superposed with the seed in Fig. 5B. However, the usefulness for sequence based search for this task is compromised by a large number of false-positive results. For example, Trace amine-associated receptor 2 (Uniprot Q9P1P5) is ranked high in the Blast results (E-value: 4.2 × 10^−49^) and has similar overall structure to the seed (Fig. 5A), but does not contain 290Phe and 312Asn amino acids which are involved in the binding of the beta-blocker drug (Fig. 5C). Act-Seek results for beta-blockers include different types of beta adrenergic (1,2 and 3), serotonine receptors and dopaminergic receptors, among others. Interaction of beta-blockers with dopaminergic receptors has been previously reported (30), yet specific receptors found by Actseek are potentially novel off-targets for beta-blockers.

**Fig. 5.**
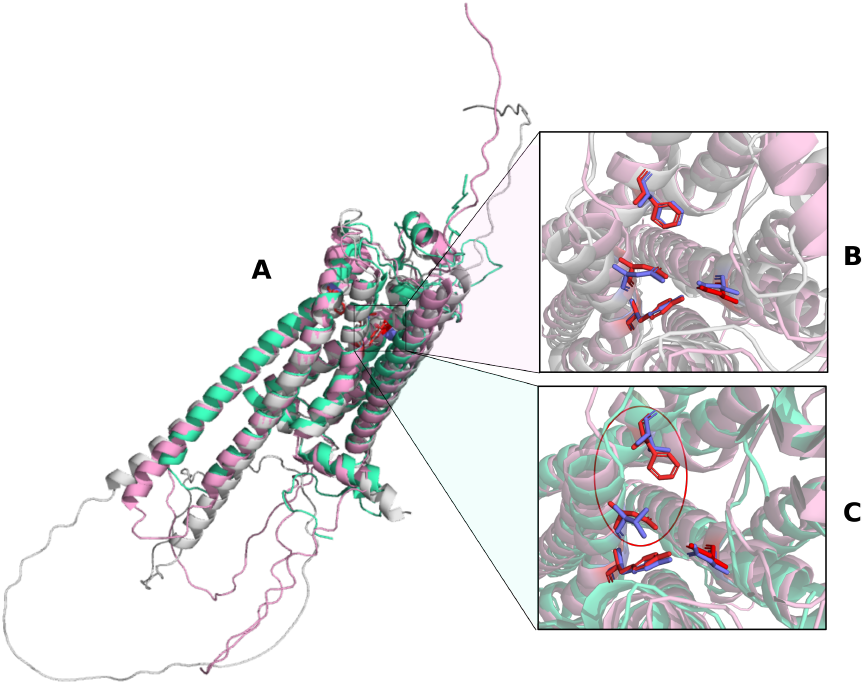
Comparison of results from ActSeek and Blast for beta-blocker off-targets. **A** Superposition of the alphafold structures for the seed (beta2 adrenergic receptor (P07550), (pink), for a protein found by both ActSeek and Blast (5-hydroxytryptamine receptor 1B (P08908), (beige), and a protein found only by Blast but not ActSeek, Trace amine-associated receptor 2 (Q96RI9), (green). **B** Superposition of active site amino acids from the seed (red), and 5-hydroxytryptamine receptor 1B (blue) found by both Actseek and Blast. **C** Superposition of active site amino acids from the seed (red) and the Trace amine-associated receptor 2 (Q96RI9)(blue), which was found by Blast but not by Actseek.

## Conclusions

We present here the ActSeek algorithm to rapidly search proteins with relevant active sites from alphafold database based on seed protein coordinates. Advantages of ActSeek to commonly used protein mining tools based on sequences (BLAST) and structures (FoldSeek) are demonstrated for finding new enzymes and drug off-targets. Our results demonstrate that ActSeek can complement searches based on sequence or overall structure by finding proteins containing the relevant active site that are not recognized based on sequence or structure. On the other hand, ActSeek can refine searches by filtering out results with similar sequence or structure, but not having the relevant active site. These features increase the probability to find new proteins with desired functions, particularly when structural details of a local active site are crucial for function. This helps to focus further screening efforts to most probable candidates.

Finding proteins with matching active site 3D conformations is particularly useful for proteins whose functions are based on specific binding certain substances, such as enzymes and drug targets. For example, ActSeek can find natural proteins from databases with similar active sites to the seed enzyme, which can be then further screened for relevant novel functionalities, such as specificity for new substrates. Expressibility and biocompatibility of such natural proteins are expected to be on average better than for *de novo* designed proteins. Another example on practical benefit is finding potential off-targets for drugs by searching human proteins with similar binding sites to known targets. Sequence based search returns many false positive results for such search, but ActSeek can filter out results that lack the relevant active sites, thereby substantially reducing the resources required for further screening.

We foresee wide range of applications for ActSeek in understanding and engineering biological systems from classification of protein functionalities for functional annotation to protein engineering, enzyme design and drug development.

## Supplementary Note 1: Material and Methods

### A. Defining the relevant amino acids in the active site

For a start, the user needs to define the binding site in the seed protein by providing cartesian coordinates or positions of the three most important amino acids. These amino acids will be used to align the query structure with the seed. User can also define additional amino acids that are relevant for the active site, which will be used for ranking when selecting the best query protein. Also neighboring amino acids in the sequence (four by default, but can be reset by the user) to the selected amino acids (binding site and additional ones) will be used for the ranking. Coordinates of C^*α*^ and C^*β*^ atoms in the amino acids will be used in the searches. User can define whether amino acid types in the local structure of the query need to exactly correspond the seed ones, or define a list of similar amino acids that are accepted at each position.

### B. Pre-screening

ActSeek algorithm first downloads the query structure from the database and checks if the three main amino acids defined by the user are present in the sequence. Before comparing local 3D configurations with the seed, ActSeek filters potential matches by comparing inter-amino acid distances. To this end, distances between the amino acids from all possible sequential combinations are calculated. Only if all inter-amino acid distances deviate less than 3Å from the values in the seed structure, the structure with the given mapping will be considered further. Notably, several possible mappings may be found for a single structure. If the number of possible combinations exceeds 2000 (can be reset by the user), this amount of combinations are randomly selected for testing.

### C. Finding the optimal rotation and translation matrices

Local 3D structures that satisfy the condition for binding site inter-amino acid distances in the pre-screening step are then further compared with the seed. This is done by finding the optimal rotation and translation matrices to overlay the binding site amino acid configuration with the seed using singular value decomposition (SVD) and least square fitting (9).

### D. Selecting the best query proteins

After performing euclidean transformations with the optimal matrices, similarities of active sites between query and seed proteins are measured by calculating distances between relevant amino acids. To this end, we first calculate the average distances for selected amino acids (binding site and additional ones) between a query structure and the seed. Additional amino acids are counted only if corresponding type is found from the query protein (exact match or correspondence defined by the user). In addition, the same distances are calculated for neighboring amino acids (set by user or 4 by default) around the each relevant amino acids. Distances between neighboring amino acids are used regardless of the types of these amino acids in the query and seed.

The score for each possible mapping of amino acids in the given structure is then calculated as an average of the distances of mapped amino acids divided by the number of amino acids over which the average is taken (to penalize situations where additional amino acids defined by the user are not found from the query). The combination of mapped amino acids with lowest score is then selected for the given structure. Finally, the given structure is selected if its lowest score was less than user defined value (default 1 Å).

### E. Other calculations. ActSeek calculates additional measures to assess the structural similarity between seed and query proteins, which can help filter the results further

- Protein alignment percentage: The alpha carbons of the query structure are aligned with those of the seed protein by optimally pairing them using dynamic programming (31). Only pairs with the distance below 2 Å are considered and used to calculate the percentage of aligned amino acids in the query structure. This value is given as a “Structural mapping percentage” in the result file.
- Local protein alignment: We calculate a weighted average of the number of 5, 10, 15, and 20 consecutive amino acid pairs that are aligned in both structures. This average is then normalized with the value from perfectly aligned structures, resulting values between 0 (local similarities not found) and 1 (structures fully aligned). This value is given by ActSeek as “Structural local similarity” in the result file. This measure indicates similarities of substructures that are aligned with the seed protein, without considering the rest of the protein that might differ significantly.
- Cavity Comparison: To compare active site cavities between query and seed proteins, cavities are first recognized using the pyKVFinder (32) algorithm. Cavities between seed and query are then compared based on above described protein alignment. We also provide the percentage of amino acids aligned in the cavity compared to the total amino acids. The result file provides these measures as ‘Cavity distances’, ‘Cavity mapping (case:seed)’, and ‘Cavity mapping percentage’. This feature is optional because it reduces the program’s performance during extensive searches.

### F. Implementation and running

The algorithm supports multiprocessing and it is designed for computer clusters, though it can run in a single computer for smaller searches. Each running node in the cluster with 24 CPUs would process 10000 proteins per 2-10 minutes. with timing dependent on the average number of possible mappings to test. ActSeek is written using python. The code can be found in https://github.com/vttresearch/ActSeek

### G. Blast searches and FoldSeek searches

We used Uniprot Knowledgebase TrEMBLE (1), containing around 253 millions of sequences as a database for the Blast search. The Evalue threshold was set to 10. We used Blastp algorithm (4) for the protein search. For the FoldSeek (7) search we used the Alphafold/UniProt50 database (3) and run it with the default parameters

### H. ActSeek search examples

#### PHA synthases

We used the PHA synthases Class I from Chromobacterium violaceum (uniprot: Q9ZHI2) as the seed, and its three catalytic amino acids 291Cys, 447Asp, and 477His (33) as the sites for the ActSeek search. Additional amino acids defined for the search were 393Trp, and 477Gly. Default values were used for other ActSeek parameters. The search was performed through Alphafold database(3) for proteins that are annotated with *αβ*-hydrolase fold in the Uniprot database (1) (total 2567710 enzymes) because all PHA synthases are expected to have such fold. We filtered the data based on local protein alignment score because we did not anticipate significant structural divergence in this case. When the mapping included only the three main catalytic amino acids (additional amino acids not found), we set the threshold for local protein alignment score to 0.4. When additional amino acids were included in the mapping, we lowered the threshold to 0.1. After filtering, we also removed any proteins that had been deleted from UniProt during the course of our study.

#### PETase

We used poly(ethylene terephthalate) (PET) depolymerase from *Piscinibacter sakaiensis* (*Ideonella sakaiensis*) as the PETase seed (Uniprot: A0A0K8P6T7), and its three catalytic amino acids 160Ser, 206Asp, and 237His (34) as the sites for the ActSeek search. Additional amino acids defined for the search were 87Tyr and 185Trp. Also phenylalanine was accepted for position 87 and tyrosine for position 185. Five neighboring amino acids around each each selected amino acids were used for the ranking (instead of default value of four). The search was performed for proteins annotated with *αβ*-hydrolase fold in the Uniprot database (1) (total 2567710 enzymes) as PETases are know to have the *αβ*-hydrolase fold (34).

#### MHETase

We used protein that catalyzes the hydrolysis of mono(2-hydroxyethyl) terephthalate (MHET) as the MHETase seed (uniprot: A0A0K8P8E7), and its amino acids 225Ser, 492Asp, and 528His as the sites for the search (35). Additional amino acids for the search were 224Cys and 529Cys. Default values were used for other ActSeek parameters. The search was performed in proteins annotated with *αβ*-hydrolase fold in the Uniprot database (1) (total 2567710 enzymes) as MHETases are know to have the *αβ*-hydrolase fold.

#### Erlotinib targets

We performed a search using the target of the drug Erlotinib as a seed (Epidermal growth factor receptor, Uniprot P00533). For the search, we used important amino acids for the binding of the drug, such as 718Leu, 790Thr, and 855Asp (36) without additional ones. Also methionine or isoleucine were accepted at position 790. After the search, we filtered the results by removing hits with the local protein alignment score lower than 0.4. We searched in all the human proteins listed in Uniprot (20,417 proteins).

#### Sorafenib targets

We performed a search using the Sorafenib target B-Raf kinase as a seed (Uniprot P15056), and its amino acids 529Thr, 532Cys and 536Ser as the sites used for the search without any additional amino acid. Also valine was accepted at position 529 and asparagine at position 536. We searched in all the human proteins listed in Uniprot (20,417 proteins).

#### Beta-blocker targets

We used the native receptor of these drugs, the beta2 adrenergic receptor (Uniprot id: P07550) as the seed. The main amino acids used for the search were 113Asp, 290Phe and 312Asn and additional ones were 114Val and 289Phe (37), which are known to be involved in the drug binding. We allowed amino acids of the same type (i.e., uncharged polar, charged, hydrophobic, and aromatic) to be interchangeable during the search. We searched in all the human proteins listed in Uniprot (20,417 proteins).

We are making the results of ActSeek, Blast and FoldSeek available in the supplementary material.

## Supplementary Note 2: Acknowledgements

We acknowledge CSC – IT Center for Science for computational resource and the Jane and Aatos Erkko Foundation (JAES) under project 220048 (Virtual laboratory for Biodesign, JAES-BIODESIGN) for their important support that allowed the realization of this work. OHSO acknowledges Research Council of Finland for financial support (358505, 356568). Additionally, we would like to acknowledge Antonio L. Rodríguez and Alejandro Revuelta for their help and advise related to the computer vision algorithms and code optimization, and Gopal Peddinti for his help running FoldSeek.

